# Multidimensional Characterization of Novel Phage UCB24 Targeting *Erwinia amylovora*

**DOI:** 10.64898/2026.06.18.733149

**Authors:** Ümran Çapar, Ömür Baysal, Ahmet Can, Kubilay K. Baştaş, Ayşegül Gur, Tuba Baygar

**Affiliations:** Molecular Microbiology Unit in Department of Molecular Biology and Genetics, Faculty of Science, Muğla Sıtkı Koçman University, 48121 Kötekli, Muğla, Turkey; Molecular Plant and Microbial Biosciences Research Unit (MPMB-RU), University of Worcester, Henwick Grove, Worcester, WR2 6AJ, United Kingdom; Department of Plant Protection Faculty of Agriculture, Selcuk University Campus/Konya/Turkey; Material Research Laboratory, Research Laboratories Center, Muğla Sıtkı Koçman University, Muğla, Turkey

**Author notes:** Corresponding author, Institutional. These authors share first authorship (Ümran Çapar, Ömür Baysal). Ümran Çapar, Ömür Baysal, Ahmet Can, Kubilay K. Baştas, Ayşegül Geduk Tuba Baygar.

## Abstract

This study evaluates the novel bacteriophage UCB24 as an eco-friendly biocontrol agent against *Erwinia amylovora*, the bacterial pathogen responsible for fire blight. Following purification, UCB24 was characterized for its optimal multiplicity of infection (MOI), infection kinetics, and environmental stability across diverse pH and temperature ranges. Microtitration assays and scanning electron microscopy (SEM) confirmed distinct morphology of the phage and its potent capacity to disrupt *E. amylovora* biofilms. Whole-genome sequencing and phylogenetic profiling identified UCB24 as a genetically distinct relative of four known phages. Furthermore, protein-protein interaction analyses revealed a strong binding affinity between the phage lysin and the host’s *N*-acetylmuramic-acid 6-phosphate etherase, uncovering the precise molecular mechanism driving targeted host destruction. *In vivo* plant trials demonstrated exceptional protective efficacy in Apple cv. Gala (88.61%) and Quince cv. Esme (81.37%), significantly outperforming traditional copper treatments under severe baseline pathogen pressure. Consequently, UCB24 represents a highly effective and sustainable biopesticide for managing fire blight in susceptible orchard ecosystems.

## 1. Introduction

*Erwinia amylovora* is a Gram-negative bacterium belonging to the family *Enterobacteriaceae* and is the causal agent of fire blight disease, which poses a global threat by infecting numerous plant species within the Rosaceae family, including apple, pear, and quince. *E. amylovora* infection spreads rapidly, particularly during the flowering period, via wind, rain, and pollen-carrying vectors; through the production of exopolysaccharides (EPS), it blocks plant vascular tissues, irreversibly disrupting water and nutrient transport (Johnson and Stockwell 1998; Yuan et al. 2022).

In addition, it exhibits an advanced pathogenicity mechanism that causes necrosis in host cells through effector proteins transferred via the Type III secretion system (T3SS), ultimately leading to plant death (Zeng et al. 2021). This disease, which causes devastating economic losses worldwide, continues to represent a strategic obstacle to Türkiye’s sustainable agriculture goals. According to the Turkish Statistical Institute (TURKSTAT) 2025 plant production projections and data, Türkiye’s total fruit production has reached approximately 29 million tons, of which around 5.5 million tons consist of pome fruits such as apple, pear, and quince that are highly susceptible to fire blight (Turkish Statistical Institute 2025). Since the 1980s, fire blight outbreaks in major fruit-producing regions of Türkiye have posed a serious threat, negatively affecting not only agricultural production but also agro-based industries and consumer markets (Öktem and Benlioğlu 1988). The disease significantly reduces fruit yield due to flower infections and, in some cases, may completely prevent fruit formation. Moreover, damage to shoots decreases the tree’s capacity to produce fruit in subsequent seasons, leading to long-term production losses. Necrotic lesions formed on branches weaken the structural integrity of the plant and may result in severe damage, progressing to the loss of entire trees in susceptible species. Nursery production areas are particularly at risk, as infected seedlings facilitate disease spread. In regions where fire blight is present, restrictions on the sale of infected seedlings and quarantine measures cause substantial economic losses for producers (Esteban-Herrero et al. 2023). Chemical control methods, such as copper-based compounds and antibiotics, are commonly used to manage *E. amylovora* infections. However, these approaches promote the development of bacterial resistance over time and pose environmental concerns (Ryu et al. 2023). In particular, prolonged and intensive use of copper-based compounds can accumulate in soil, adversely affecting plant growth and causing harmful ecological consequences. Furthermore, existing chemical and cultural control strategies have proven insufficient to completely eradicate the disease (Park et al. 2017). Therefore, environmentally friendly and sustainable alternative control strategies are urgently needed. In this context, bacteriophages have emerged as a promising option for the biological control of plant pathogens. Bacteriophages are viruses that infect bacteria and are naturally found wherever bacteria exist (Dion et al. 2020). Due to their host specificity and their inability to induce antibiotic resistance, bacteriophages are considered effective and environmentally sensitive biological agents for controlling plant diseases (Pinheiro et al. 2020; Greer et al. 2025).

Within the scope of this study, the characterization of our bacteriophage (UCB24) effective against *E. amylovora* and the investigation of its biocontrol potential were aimed. Initially, our isolated bacteriophages were purified and subsequently subjected to enrichment procedures. Genomic DNA was isolated from phage concentrates for genomic analyses. Following whole-genome sequencing, the obtained DNA sequence was analyzed and digested with selected restriction enzymes to generate restriction enzyme profiles. Subsequently, for phage characterization, the multiplicity of infection (MOI) was determined; phage activity under different pH and temperature conditions was evaluated; killing curve analysis was performed; and the effects on *E. amylovora* biofilm were investigated using a microtitration method. In addition, the isolated bacteriophage was characterized at both genomic and microscopic levels. The morphological structure of bacteriophage particles was visualized using scanning electron microscopy (SEM), allowing the analysis of both phage structural features and the effects of phage application on *E. amylovora* biofilm. The disruptive effect of phage treatment on biofilm structure was considered one of the key findings supporting the biocontrol potential of the phage. At the genomic level, phage DNA was isolated, sequenced, assembled, and annotated, resulting in the mapping of the phage genome. Furthermore, phylogenetic analyses were conducted to determine the phage’s closest relatives. The results were subjected to statistical analysis and demonstrated in vitro assays that the isolated phage possesses strong biocontrol potential against *E. amylovora*. Accordingly, bacteriophages are regarded as an alternative and environmentally friendly approach for biological control, and this study provides a scientific basis for future applied and field-based research.

## 2. Materials and method

### 2.1 Bacterial strains and culture conditions

The bacterial strain *E. amylovora*, used as the host strain for bacteriophage detection, was obtained from Department of Plant Protection at Selçuk University, Konya-Türkiye. The bacterial strain was cultured on Nutrient Agar (NA) medium and incubated at 28 °C for 18 h. Subsequently, a single colony grown on solid medium was transferred into a Falcon tube containing Nutrient Broth (NB) and cultured in a shaking incubator at 150 rpm and 28 °C for 16 h. The resulting culture was used in subsequent experiments (Vique et al. 2025).

### 2.2 Bacteriophage isolation and purification

To isolate bacteriophages (UCB24) specific to *E. amylovora*, soil samples were collected beneath apple, pear, and quince trees in the Samandağ district of Hatay province, which were suspected to exhibit fire blight symptoms. The collected samples were transferred into sterile Falcon tubes, NB was added to a final volume of 20 mL, and the mixtures were homogenized using a vortex. The samples were incubated in a shaking incubator at 28 °C and 250 rpm for 2 h. Following incubation, the samples were centrifuged at 2,000×g for 10 min to allow soil particles to settle, and the supernatant was filtered through a 0.22 µm pore-size filter to remove cellular debris.

Initial detection of phage presence was performed using a spot test with the double-layer soft agar method. Briefly, 200 µL of *E. amylovora* (strain EAKkb29) culture was mixed with 5 mL of soft agar (0.8% NB, 0.7% agar) and poured onto NA plates as a thin overlay. After the agar solidified, 5 µL of the filtered samples was spotted onto the plate surface, and the plates were incubated at 28 °C for 18 h. The appearance of lysis zones was considered indicative of phage activity (Wang et al. 2022).

For purification of phage particles from positive samples, the double-layer agar method was employed. A total of 500 µL of lysis-positive filtrate was mixed with 500 µL of *E. amylovora* culture and 5 mL of soft agar, then poured onto NA plates. After overnight incubation at 28 °C, well-defined and transparent phage plaques were selected and collected using sterile pipette tips, then suspended in 100 µL of SM buffer [10 mM Tris-HCl (pH 7.5), 10 mM MgSO-1, 68 mM NaCl, 1 mM CaCl-1]. The obtained phage suspensions were serially diluted, and spot tests were repeated to verify each dilution step (Sabri et al. 2022).

### 2.3 Host range determination

The host range of the isolated bacteriophage was determined by spot testing against different bacterial strains available in the laboratory culture collection, including *Clavibacter michiganensis* ssp. *michiganensis*, *Pseudomonas syringae* pv. *tomato*, *Xanthomonas vesicatoria*, *Klebsiella pneumoniae, Klebsiella oxytoca, and Bacillus subtilis*. Each bacterial strain was grown overnight in NB medium at its appropriate incubation temperature. A volume of 100 µL of bacterial culture was added to cooled soft agar and spread onto NA plates. Ten microliters of the purified phage suspension were spotted onto the plates, which were then incubated overnight at suitable temperatures for each bacterium to observe plaque formation (Sabri et al. 2022).

### 2.4 Stability test of bacteriophage under various conditions

The stability of phages (10-1 PFU/mL) at different temperatures (15, 25, 30, 37, and 45 °C) was evaluated after incubation in a water bath for 1 h (Lim et al. 2013). Similarly, the pH stability of phages (10-1 PFU/mL) at different pH values (5, 6, 7, 8, and 9) was assessed following incubation at 28 °C for 1 h. All experiments were conducted in triplicate, and phage titers were determined using the double-layer agar plating method (Li et al. 2022).

### 2.5 Optimal moi and killing curve assay

Multiplicity of infection (MOI) assays were conducted to determine the ratio of bacteriophages capable of infecting potential host cells simultaneously. *E. amylovora* cultures (10-1 CFU/mL) grown in NB were mixed with phages at different MOI values (100, 10, 1, and 0.1 PFU/CFU). After a 10 min adsorption period, free phages were removed by centrifugation at 15,000 g for 5 min at 4 °C, and the pellet was resuspended in 5 mL NB and incubated again at 27 °C for 4 h. Following incubation, the culture was centrifuged at 15,000 g for 5 min at 4 °C, and the supernatant was filtered through a sterile 0.22 µm syringe filter. Phage titers were then determined using the double-layer agar method, and the MOI yielding the highest phage production was considered the optimal MOI (Lee et al. 2021).

To assess the effect of the isolated bacteriophage on bacterial growth, a killing curve assay was performed. Briefly, 40 µL of *E. amylovora* culture (10-1 CFU/mL) and 40 µL of fresh NB were added to each well of a 96-well microplate. The bacteria were then infected with 40 µL of phage suspension at different MOI values (0.01, 0.1, 1, 10, and 100). The plate was incubated at 27 °C, and optical density was measured at 600 nm every hour for 12 h. All experiments were conducted in triplicate (Jamal et al. 2015).

### 2.6 Biofilm inhibition assay

The formation of *E. amylovora* biofilm and the inhibitory effects of bacteriophages on biofilm formation were evaluated using a microtiter plate assay. For this purpose, 200 µL of *E. amylovora* culture grown in NB supplemented with 2% glucose was inoculated into each well of a 96-well microplate. Distilled water was added to control wells as a negative control. After incubation at 27 °C for 24 h, the contents of the wells were aspirated and removed. To assess the effect of phages on biofilms, 200 µL of phage suspension was added to each well, and the plates were incubated under the same conditions for an additional 24 h. At the end of incubation, both phage-treated and control wells were washed three times with phosphate-buffered saline (PBS) and fixed with 99% methanol for 15 min. After air-drying, the biofilms were stained with 1% crystal violet, and excess stain was removed with distilled water. The stained biofilm was solubilized with 33% glacial acetic acid, and absorbance was measured at 570 nm to quantify biofilm biomass (Liu et al. 2025).

### 2.7 SEM-based visualization of biofilm disruption and phage morphology

Scanning electron microscopy (SEM) was used to evaluate the effects of phage treatment on *E. amylovora* biofilm and to visualize bacteriophage particle morphology. Sterilized glass coverslips were placed in sterile Falcon tubes containing 5 mL NB, and each tube was inoculated with bacterial suspension at a concentration of 1×10-1 CFU/mL. Phage-treated and untreated groups were prepared separately and incubated for 24 h. After incubation, the coverslips were gently washed with PBS to remove unattached cells, fixed with 2.5% glutaraldehyde for 4 h, washed again with PBS, and dehydrated through a graded ethanol series (50%, 70%, 90%, and 96%). The samples were air-dried, gold-coated, and examined using a scanning electron microscope (Jeol JSM-7600F, Japan) at an accelerating voltage of 15 kV (Yuan et al. 2019).

For morphological visualization of bacteriophage particles, 1.5 mL of phage suspension (phage + SM buffer) was fixed with 2.5% glutaraldehyde for 4 h, centrifuged at 18,000 g for 120 min, and the pellet was gently washed with PBS. The phage particles were dehydrated through graded ethanol concentrations (50%, 70%, 90%, and 96%), air-dried on sterile coverslips, gold-coated, and examined by SEM under the same conditions (Phee et al. 2013).

### 2.8 Genome sequencing and bioinformatic analysis

Genomic DNA was isolated from the bacteriophage for genomic analyses. Briefly, 2 mL of filtered phage suspension was centrifuged at 15,000 rpm for 2 h at 4 °C, and the resulting phage pellet was resuspended in 100 µL SM buffer. To eliminate bacterial DNA contamination, DNase treatment was performed at 37 °C for 30 min, followed by DNase inactivation at 70 °C. Proteinase K treatment was applied at 55 °C for 1 h to remove proteins. DNA extraction was then carried out using phenol–chloroform–isoamyl alcohol, followed by precipitation with isopropanol/sodium acetate according to Sambrook (2001). The DNA pellet was washed with 70% ethanol, air-dried, resuspended in 50 µL TE buffer, and stored at −20 °C. The purified DNA was used to construct a library with blunt triple junctions consisting of fragments of approximately 500 base pairs in length. Subsequently, the library was subjected to sequencing using an Illumina Hi-Seq 2500 sequencing system (Refgen Biotechnology Ltd. Ankara, Türkiye) for whole-genome sequencing using the Illumina HiSeq 2500 platform with ∼500 bp fragment libraries (Kim et al. 2025).

Raw sequencing reads in FASTQ format were subjected to initial quality assessment using Falco (Galaxy Version 1.2.4). To ensure high-quality data for assembly, adapters and low-quality bases (Phred score < 30) were trimmed using fastp, with a minimum length requirement of 50 bp. To eliminate potential host DNA contamination, reads were competitively mapped against the reference genome of the bacterial host (*E. amylovora* CFBP1430) using Bowtie2. Only unmapped reads representing the enriched viral fraction were retained for downstream analysis. De novo assembly of the filtered reads was performed using the SPAdes (v3.9.0.) assembler with the meta flag to account for potential variations in viral coverage. For long-read datasets, Flye was utilized in metagenomic mode. The resulting contigs were evaluated for circularity by identifying terminal redundancies using CheckV, which also provided estimates of genome completeness and contamination levels. Protein-coding sequences (CDS) were predicted using Pharokka (rapid standardised annotation tool for bacteriophage genomes and metagenome tool (Galaxy Version 1.3.2+galaxy) with phanotate and prodigal option as it is optimized for the dense and overlapping gene structures characteristic of bacteriophage genomes. Functional annotation was assigned by searching predicted amino acid sequences against the NCBI NR databases using HMMER and BLASTp (E-value cutoff, 10^-5^), respectively. Open reading frames (ORFs) were predicted using ORF finder (NCBI). Gene identification was further supported using PHASTER and NCBI BLASTn. Specialized motifs, such as tRNA genes, were identified using tRNAscan-SE. For taxonomic classification, a proteomic tree was constructed using the VICTOR (Virus Classification and Tree Building Online Resource) pipeline. Particular emphasis was placed on identifying the viral lineage based on the RNA polymerase-encoding gene. The obtained whole sequence data was submitted to the NCBI database for GenBank.

The entire analysis was also carried out by the VICTOR web service (https://victor.dsmz.de), a method for the genome-based phylogeny and classification of prokaryotic viruses (Meier-Kolthoff and Göker 2017). The resulting intergenomic distances were used to infer a balanced minimum evolution tree with branch support via FASTME including SPR postprocessing (Lefort et al. 2015) Taxon boundaries at the species, genus and family level were estimated with the OPTSIL program (Göker et al. 2009), the recommended clustering thresholds (Meier-Kolthoff and Göker 2017) and an F value (fraction of links required for cluster fusion) of 0.5 (Meier-Kolthoff et al. 2014).

### 2.9 Molecular fingerprinting of phage

Genomic DNA of the isolated phage was subjected to *in silico* restriction analysis using whole-genome sequencing data. Restriction enzyme digestion patterns were predicted, and BamHI, HinfI, and PfeI were selected based on the analysis. Restriction digestions were then performed on phage genomic DNA, and the resulting fragments were separated on 1% agarose gel electrophoresis. The banding patterns obtained from gel electrophoresis were compared with *in silico* predictions (Karthika et al. 2021).

### 2.10 Genomic integration analysis between phage and host *E. amylovora*

Genomic sequences for the host bacterium, *E. amylovora* (Accession Nr. CFBP1430), and the novel bacteriophage (Accession: PV930069) were processed in the R statistical environment (v4.5.2). Sequences were managed using the Biostrings package. To ensure high-quality alignment, the phage assembly was filtered to remove contigs shorter than 200 bp and verified for sequence integrity prior to comparative analysis.

Large-scale genomic synteny was determined using the DECIPHER package. A local SQLite database was initialized to facilitate the comparison. Sequences were loaded into the database using the Seqs2DB function with unique identifiers. Syntenic blocks were identified via the FindSynteny algorithm, which utilizes maximal exact matches (MEMs) to define regions of homology. The resulting synteny map was visualized to identify genomic rearrangements, inversions, and conserved modules between the phage and the bacterial chromosome.

To characterize the “Interaction Landscape” between the phage and the host, a sliding-window scanning approach was implemented. The E. amylovora genome was partitioned into 100 equidistant bins (approx. 38 kb per bin). A 50 bp genomic probe, corresponding to the predicted phage integration signal (including the 34 bp attP site), was scanned across the host genome using the matchPattern function. The interaction potential was quantified by match density per bin, allowing for a maximum of 12–15 mismatches to account for natural genetic drift and divergent homologous regions. The resulting density matrix was visualized as a spatial heatmap using the ggplot2 and pheatmap libraries. A color scale was applied to distinguish between genomic structural preservation and putative integration sites. For the interaction heatmap, hierarchical clustering was disabled (cluster_rows = FALSE) to maintain the linear coordinate integrity of the bacterial chromosome. The final analytical pipeline was documented in R scripts to ensure computational reproducibility convenient for standards on GitHub.

### 2.11 Protein-protein interaction analysis

To investigate the interaction between N-acetylmuramic-acid 6-phosphate etherase (*E. amylovora*) and Phage lysin (EC 3.2.1.17) (Phage muramidase), we conducted computational analysis to validate the presence of unique sequence belonging to UCB24 annotation data and reference *E. amylovora* CFBP1430. We retrieved the sequences and translated to amino acid sequences using ExPasy tool and resulting output was submitted to Swiss MODEL for selecting the best matching model with high score (Gasteiger et al. 2003; Waterhouse et al. 2018). These data served as a base for modelling the corresponding, which exhibited similarity with the active sites of N-acetylmuramic-acid 6-phosphate etherase and Phage muramidase as templates for modelling SWISS MODEL (http://swissmodel.expasy.org) was used considering Z-score. Protein-protein docking was performed using the HADDOCK server (Table 2) (Van Zundert et al. 2016). The active site/interface residues of both proteins were given in Online Resource 1. The interacting residues were visualized using HADDOCK, PyMol and LigPlot+ (Schrödinger and DeLano 2023). The interaction was visualized using PyMOL Molecular Graphics System (ver. 3.1.3), integrated with Python version 3.12.4.

### 2.12 Phage efficiency against target pathogen and *in vivo* assay

The plant material used in the experiments was commercially obtained from Fodul Nursery, a producer of soft-seeded fruit saplings in Konya province. The apple cv. Gala, pear cv. Santa Maria and quince cv. Esme saplings used in the experiments were selected from healthy saplings showing homogeneous development at the age of 3. The plants were planted in 3 kg pots, and the growing medium was prepared with peat, well-rotted animal manure, and fertile field soil (a 1:1:1 mixture). During cultivation and experiments, the plants were kept in 16 hours of light and 8 hours of darkness, with 65-70% relative humidity and 26-28°C. After bacterial inoculations, the plants were additionally moistened by misting for 2 days.

The prepared UCB24 phage suspension was applied to the leaves and stems by spraying at a dose of 1x10^8^ PFU ml^-1^ using a pressurized hand sprayer. Applications were made when the annual fresh shoots of 3 different host plants (apple cv. Gala, pear cv. Santa Maria and quince cv. Esme plants) reached 25-30 cm, ensuring that all shoots and leaves were wetted. Three days after the first application, *E. amylovora* inoculation was performed, and then a second phage application was repeated 3 days later as described above (Boulé et al. 2011).

*E. amylovora* str. EAKkb21, has 98% virulence, stated in the culture collection of the Molecular Plant Bacteriology Laboratory, Department of Plant Protection, Faculty of Agriculture, Selçuk University, was used in the experiments. Stock cultures of the pathogen were inoculated onto Nutrient Agar (NA) medium, and fresh cultures after 48 hours were prepared at a concentration of 1x10^8^ CFU ml^-1^ (OD: 600, 0.15, Eppendorf Biophotometer) and inoculated. After adjusting the concentration of the bacterial suspension, the strain was kept on ice until inoculation. Using sterile scissors dipped in the prepared bacterial suspension, the tips of the young leaves of the plant were cut off and cutted leaves immersed in the suspension for about 30 seconds (Bonasera et al. 2006).

A total of nine shoots with leaves were inoculated from each host, with 3 seedlings and at least 3 fresh shoots/leaves from each plant being inoculated. In the experiments, copper sulfate equivalent to 65.82 g/L metallic copper was used as a control treatment at a dose of 125 ml/100 L water. After disease symptoms ceased, disease severity (%) was assessed by measuring the length of necrotic tissues approximately 2 months after inoculation, according to Aldwinckle et al. (2002) and Norelli et al. (1988). In the case where shoot blight did not occur, leaf symptoms were evaluated using a 0-10 scale, seven days after inoculation (Norelli et al. 1988).

The effectiveness of phage applications was determined according to Abbott (1925), and the formula is given below:

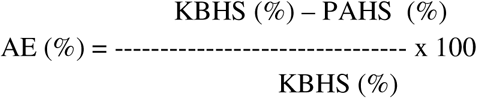

**AE:** Effectiveness of the application

**KBHS:** Average disease severity in control plants

**PAHS:** Average disease severity in phage-treated plants

In accordance with Koch’s postulates, the pathogen was re-isolated from plants showing disease symptoms (Baysal and Zeller 2004) and the purified bacterial isolates were re-identified by biochemical, physiological and molecular tests. Accordingly, Gram reaction, growth on King’s B and Crosse and Goodman media, growth at 37°C, levan formation, catalase, oxidative-fermentative activity, esculin hydrolysis, gelatin liquefaction, pectolytic activity, and acid production from erythritol and sorbitol were tested for the re-isolated and purified bacteria. DNA isolation from the re-isolated bacteria was performed using the QIAamp DNA Mini Kit (Qiagen, Hilden, Germany). The quantity and purity of the obtained DNA were evaluated by measuring absorbance at 260 nm and 280 nm wavelengths using µpower and an Eppendorf Biophotometer Plus (Eppendorf AG, Hamburg, Germany). Specific A/B primers and a PCR protocol were used for the identification of the pathogen by PCR (Bereswill et al. 1992). In the statistical analysis of the obtained data, analysis of variance was performed using MINITAB version 18. Tukey’s multiple comparison test, implemented in the MSTAT program, was used for the analysis of the data (Düzgüneş et al. 1987).

## 3. Results

### 3.1. Bacteriophage isolation and purification

For the isolation of *E. amylovora* bacteriophages, soil samples were collected from beneath pear trees exhibiting symptoms of fire blight. The collected samples were processed into suspension form, and the presence of phages capable of lysing *E. amylovora* was detected using the spot test method (Fig. 1a). To enable purification, suspensions from phage-positive soil samples were mixed with bacterial culture and soft agar and applied to NA plates using the double-layer agar method. Following incubation, bacteriophages were isolated from lysis plaques using the plaque-picking method (Fig. 1b). Selected transparent phage plaques were suspended in 100 µL of SM buffer, and the resulting phage suspensions were serially diluted. To obtain a pure single phage population, this procedure was repeated three times. To verify the dilution steps, spot tests were performed by spotting 5 µL of dilutions ranging from 1 to 10-1-1 onto double-layer agar plates prepared with *E. amylovora*; SM buffer was used as a control. Following incubation, lysis zones corresponding to the dilutions were observed, confirming successful purification (Fig. 1c).

**Fig. 1.**
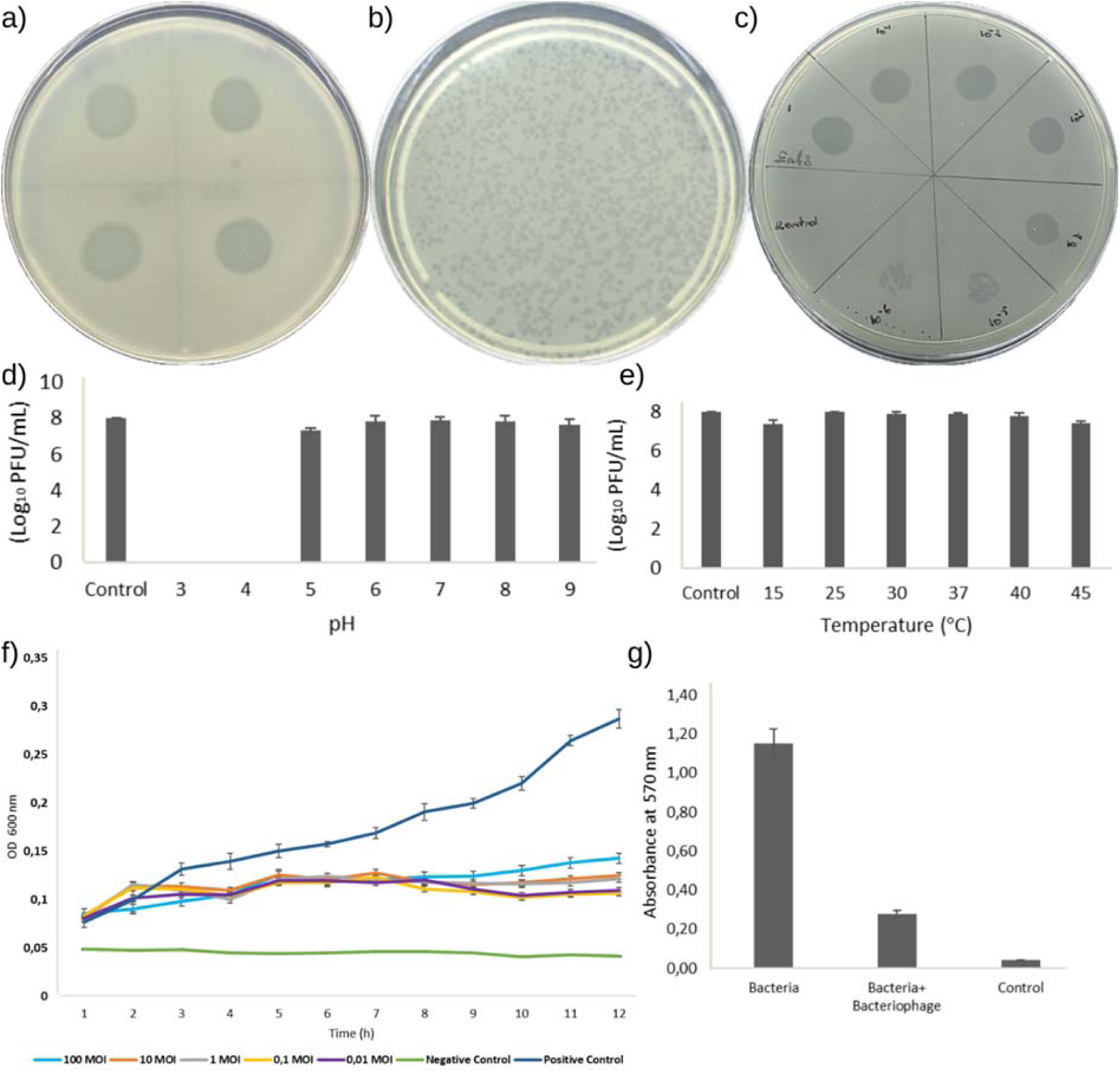
Isolation and characterization of *E. amylovora*-specific bacteriophage. (a) Phage plaque morphology obtained using the double-layer agar method; (b) lytic activity on the host bacterial lawn; (c) phage titration by spot test method using serial dilutions from 1 to 10LJLJ (SM buffer was used as control); (d) stability of the bacteriophage at different pH values (3–9); (e) thermal stability of the phage at different temperatures; (f) time-dependent in vitro bacterial growth inhibition curves at different multiplicities of infection (MOI 0.01–100) (error bars represent ± SD; statistical differences were determined using the LSD test (p < 0.05)); (g) inhibitory effect on biofilm formation determined by the crystal violet method. Data represent the mean of three independent experiments (error bars ± SD; statistical differences were determined using a one-sample t-test(p<0.05))

### 3.2. Host range of the isolated phage

Following isolation, the host range of the phage was determined using the spot test method. Phage suspensions were spotted onto plates inoculated with different bacterial strains and incubated for 18 h at temperatures appropriate for each bacterium. After incubation, lysis zones were evaluated based on plaque morphology. While the phage formed lysis plaques *on E. amylovora*, no lytic activity was observed against *Clavibacter michiganensis subs. michiganensis*, *Pseudomonas syringae pv. tomato*, *Xanthomonas vesicatoria*, *Klebsiella pneumoniae*, *Klebsiella oxytoca*, or *Bacillus subtilis*.

### 3.3. Stability test of bacteriophage

The stability of the phage (10-1 PFU/mL) under different temperature conditions (15, 25, 30, 37, and 45 °C) and pH values (3, 4, 5, 6, 7, 8, and 9) was evaluated by determining phage titers following plating. Phage stability was assessed across a broad range of pH and temperature conditions. The phage produced lysis plaques at pH values of 5, 6, 7, 8, and 9, whereas no lytic activity was detected at pH 3 and 4 (Fig. 1d). In addition, the phage exhibited lytic activity at all tested temperature conditions (Fig. 1e).

### 3.4. Optimal moi and killing curve assay

Multiplicity of infection (MOI) assays were performed to determine the ratio of bacteriophages capable of simultaneously infecting potential host cells. MOI values ranging from 0.01 to 100 were tested for E. amylovora, and the optimal MOI was determined to be 0.1.

A killing curve assay was conducted to evaluate the effect of the isolated bacteriophage on bacterial growth. Bacterial cultures were infected with phages at various titers and incubated for 12 h, and reductions in bacterial growth were assessed by measuring optical density. Changes in optical density (OD600) were compared with those of the phage-free control culture. All tested phage titers resulted in a reduction in bacterial growth. During the first two hours of incubation, no visible difference was observed between the bacterial control and the experimental groups (phage + bacteria). From the third hour onward, a significant decrease in bacterial growth was observed in the experimental groups compared with the control. All evaluated phage titers showed reduced optical density at 600 nm relative to the control. The greatest reduction was observed at MOI values of 0.1 and 0.01 (Fig. 1f).

Killing curve analysis and biofilm inhibition assays conducted to evaluate the interaction between the bacteriophage and *E. amylovora* statistically demonstrated that the phage exerted a significant inhibitory effect on bacterial growth. In vitro killing curve experiments showed a significant difference starting from the 8th hour of incubation, at which point the suppressive effect of the bacteriophage on bacterial growth became evident compared with the control. This effect remained statistically significant after the 8th hour (Supplementary Data File 1). In addition, biofilm inhibition assay results exhibited statistically significant differences compared with the control group (Supplementary Data File 2).

### 3.5. Biofilm Inhibition Assay

Biofilm production by *E. amylovora* isolates and the ability of the bacteriophage to inhibit and eliminate bacterial biofilms were evaluated using the microtiter plate assay after 24 h of incubation. Total biofilm biomass was quantified using the crystal violet staining method. According to the results, the phage significantly reduced biofilm formation by *E. amylovora*. Repeated experiments demonstrated that the phage reduced bacterial biofilm formation by an average of 4.10-fold (Fig. 1g).

### 3.6. SEM-Based visualization of biofilm disruption and phage morphology

Scanning electron microscopy (SEM) analyses revealed marked differences in biofilm formation and structure between experimental groups. In the control group containing only *E. amylovora*, a dense biofilm matrix was observed on the surface (Fig. 2a, 2b). In contrast, the phage-treated group exhibited substantial disruption of the biofilm structure, loss of bacterial cell integrity, and the presence of numerous cellular debris and lysed bacterial remnants on the surface (Fig. 2c, 2d).

**Fig. 2.**
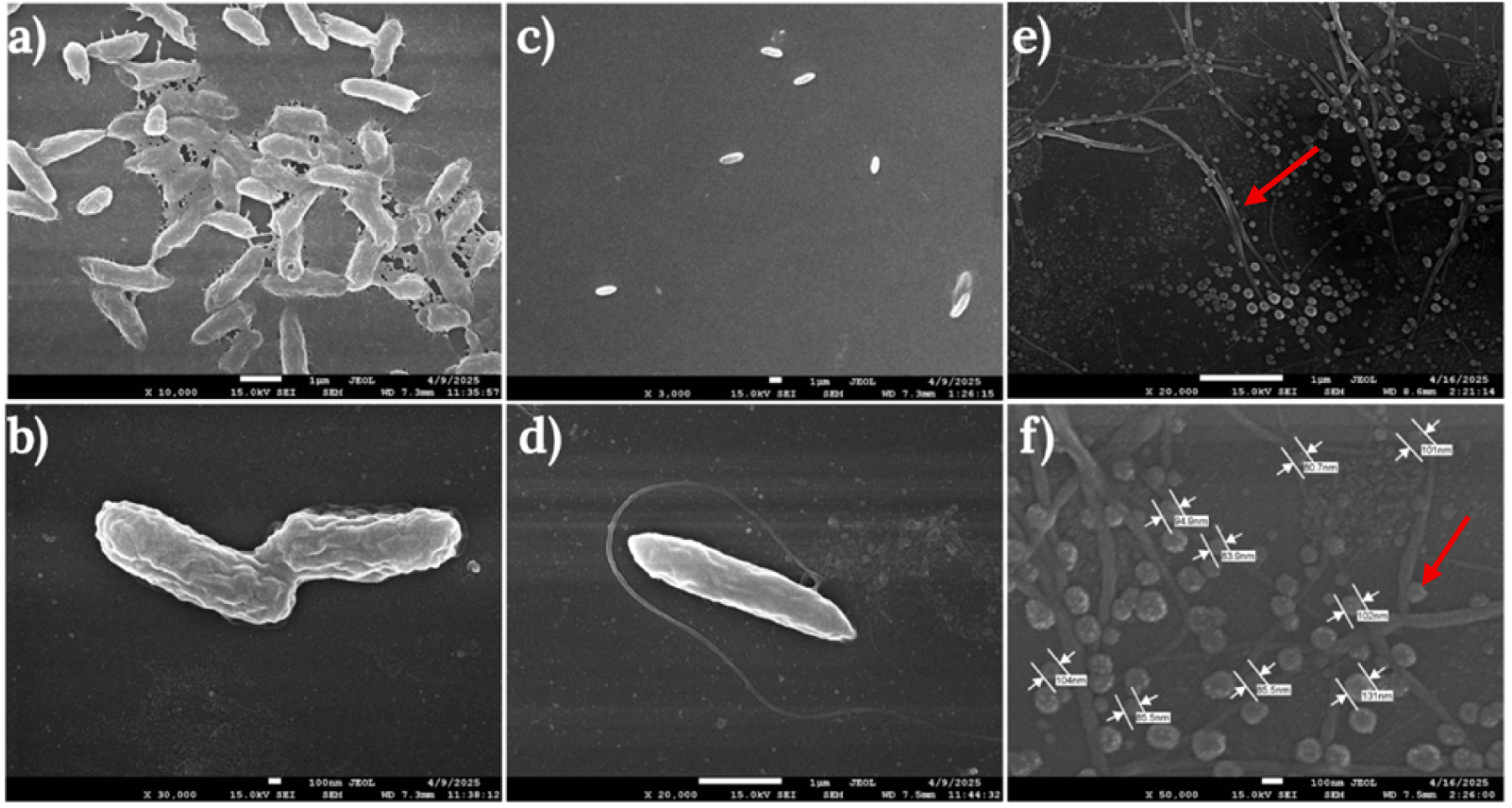
SEM visualization of host bacterial biofilm and bacteriophage particles. (a, b) Control biofilm of *E. amylovora* after 24 h incubation; (c, d) biofilm of *E. amylovora* following bacteriophage treatment over the same time period; (e, f) SEM images of the bacteriophage particles.

The lytic bacteriophage specific to *E. amylovora* was visualized by SEM, and the phage was determined to possess a capsid approximately 96 nm in diameter. In the group containing only phage particles, no biofilm formation or bacterial presence was observed (Fig. 2e, 2f).

### 3.7. Genomic characterization of UCB24

Open reading frames (ORFs) were predicted, and genome annotation was conducted using the RAST Rapid Annotation system via the Subsystem Technology server (https://rast.nmpdr.org). Gene identification was further supported using RASTK server and NCBI BLASTn. The genomic map of the bacteriophage is presented in Figure 3. The obtained sequence data were also submitted to the NCBI database under accession number PV30069.

**Fig. 3.**
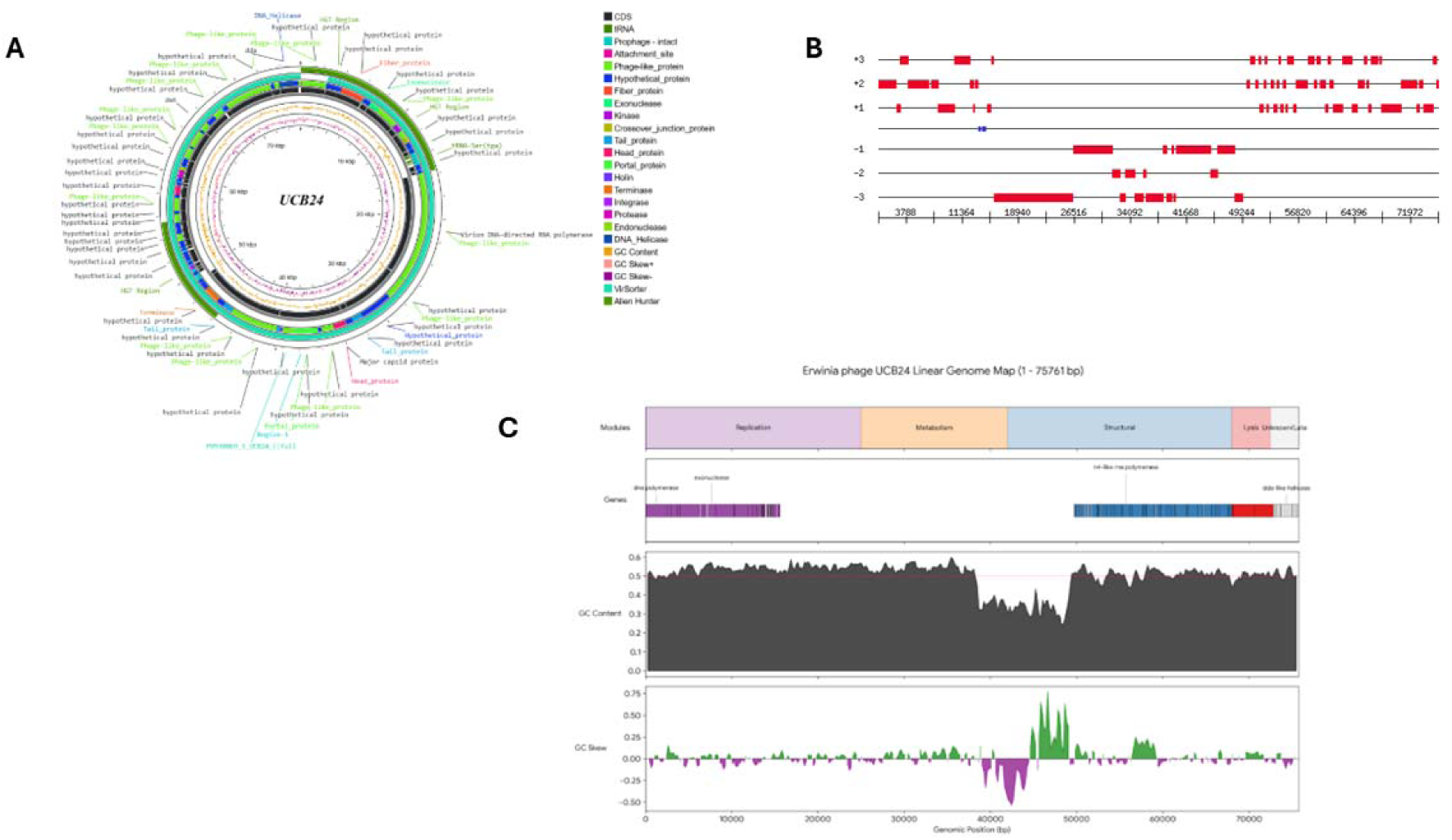
Comprehensive genomic organization and sequence analysis of the novel *E.* amylovora phage UCB24. a) Circular genome map of UCB24 visualizes the complete circular genome of the phage. The outermost rings display the predicted open reading frames (ORFs) and coding sequences (CDS), which are color-coded based on their assigned functional categories (e.g., structural proteins, replication/metabolism, lysis machinery, and hypothetical proteins) as detailed in the accompanying key. The inner concentric tracks represent the local GC content (black) and GC skew (purple/green), highlighting variations in nucleotide composition across the structural and functional boundaries of the genome. b) Linear open reading frame (ORF) distribution. The physical mapping of predicted genes across all six possible reading frames (+1, +2, +3 on the forward strand; -1, -2, -3 on the reverse strand). The red blocks represent the positioning and length of individual ORFs plotted against the genomic coordinates (spanning from 1 to 75,761 base pairs), demonstrating the coding density and strand transcription orientation of the phage. c) Linear modular map and nucleotide composition profiling. This multi-panel linear map breaks down the genome into functional and compositional segments along the horizontal genomic axis (bp): The genome is partitioned into clearly defined functional modules as replication (purple), metabolism (orange), structural (blue), lysis (red), and unknown/late elements (grey). The corresponding genes (e.g., DNA polymerase, DNA helicase) are mapped directly below their respective modules. A line plot charting the local GC percentage. A notable drop in GC content is visible near the ∼35-45 kb region, corresponding to the transition between the metabolism and structural modules. A bar plot measuring the relative abundance of guanine over cytosine (green indicates positive skew, purple indicates negative skew). Distinct shifts in GC skew often correlate with the origin and terminus of replication or demarcate major transcriptional shifts between gene cassettes.

**Table 1.** Key feature of UCB24 genome according to genomic range.

| Module | Genomic range (bp) | Key features |
| --- | --- | --- |
| Replication | 1 – 25.000 | DNA Polymerase, Helicase, Primase, and SSBs. |
| Metabolism | 25,000 – 42.000 | Nucleotide metabolism (Ribonucleotide Reductases). |
| Structural | 42.000 – 68.000 | Capsid, Portal, Tail Tape Measure, and Tail Fibers. |
| Lysis | 68.000 – 72.500 | Holin and Endolysin (Cell wall degradation). |
| Unknown/late | 72.500 – 75.761 | Hypothetical proteins and regulatory elements |

In addition to genomic analysis, we have also supported our finding with fingerprinting analysis on our phage DNA.

### 3.8. Fingerprinting and genomic comparison

The complete genome sequence of the isolated phage was subjected to in silico restriction analysis using NEBcutter software (https://nc3.neb.com/NEBcutter/). Restriction patterns generated by different enzymes were evaluated, and BamHI, HinfI, and PfeI were selected for experimental validation. Restriction digestion of phage genomic DNA was performed using these enzymes, and the resulting fragments were resolved by 1% agarose gel electrophoresis. The observed banding patterns were consistent with in silico predictions, providing preliminary insights into the restriction enzyme profile of the phage (Fig. 5).

**Fig. 4.**
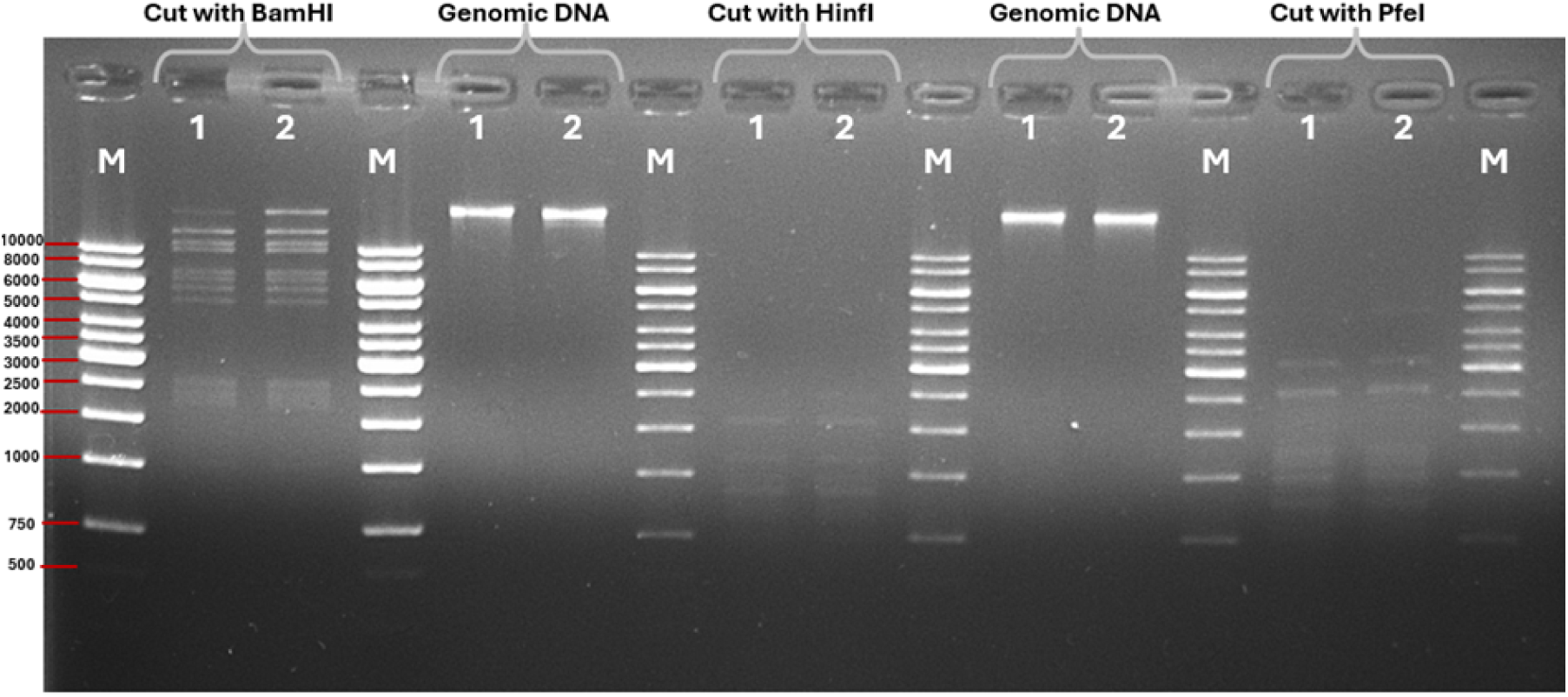
Restriction enzyme profiles of bacteriophage DNA digested with BamHI, HinfI, and PfeI. M: 1 kb DNA Ladder (Thermo Scientific).

**Fig. 5.**
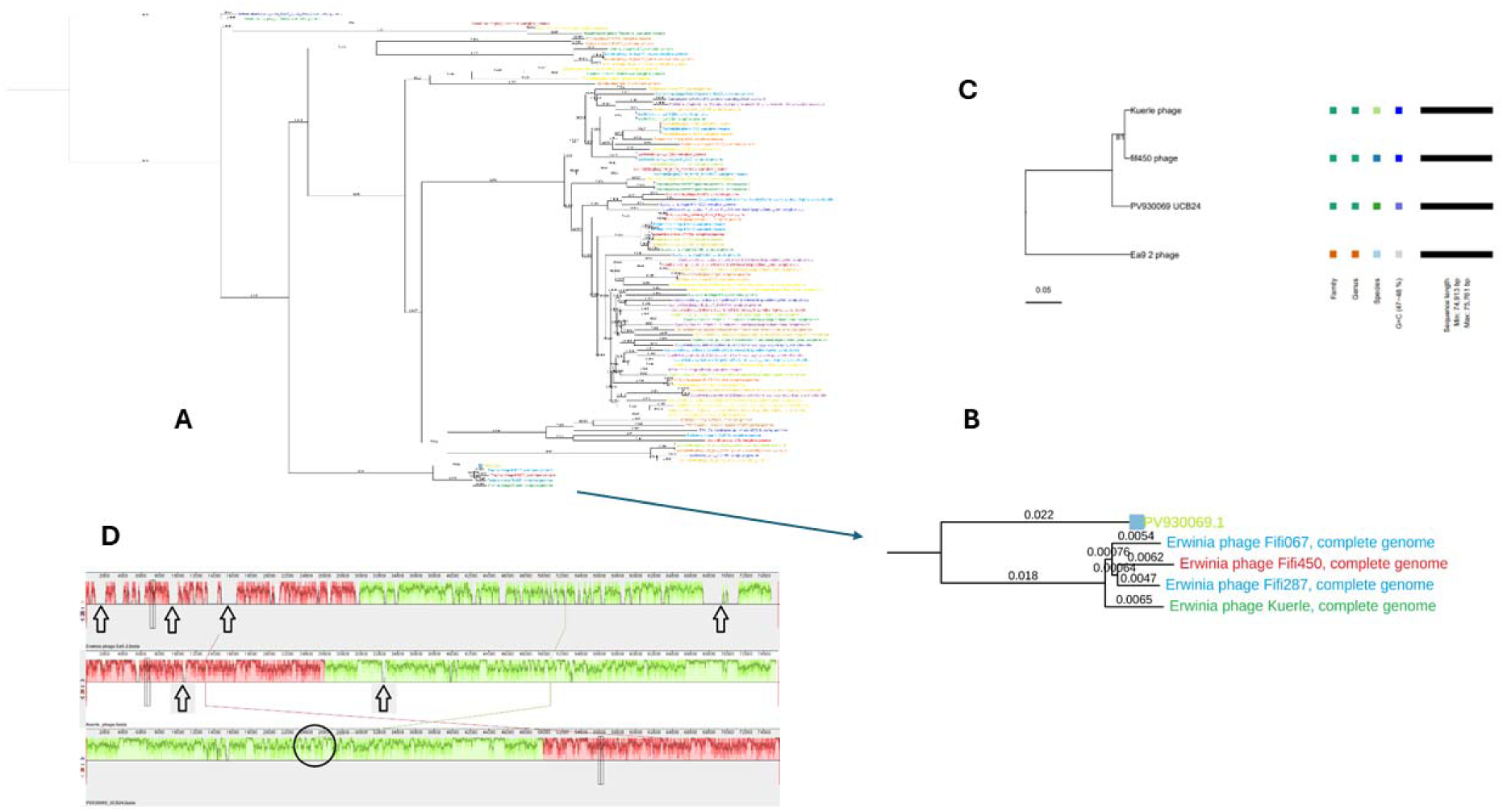
Phylogenetic and comparative genomic analysis of *Erwinia* phage UCB24. a) Whole-Genome Phylogenetic Tree: A comprehensive phylogenetic tree constructed based on whole-genome sequences utilizing the NCBI Blast tool. The evolutionary relationships were inferred using the Tamura–Nei genetic distance model under the Neighbor-Joining tree construction method. b) Clade Analysis. An expanded, detailed view of the specific clade from panel A (indicated by the blue arrow) containing the novel phage UCB24 (labeled as PV930069.1). This subfigure demonstrates that UCB24’s closest relatives form a group involving the *Erwinia* phages Fifi067, Fifi450, Fifi287, and Kuerle. c) Phylogenomic GBDP Tree: A tree inferred using VICTORnt tree trimming, which yielded an average support of 81%. The numbers positioned above the branches represent GBDP pseudo-bootstrap support values derived from 100 replications. The branch lengths are scaled in terms of the respective distance formula used. The colored squares to the right estimate taxon boundaries, plotting the phage’s classification at the family, genus, and species levels. d) Whole-Genome Alignment: A collinearity map comparing three phage genomes: *Erwinia* phage Ea9-2 (Top), Kuerle phage (Middle), and the novel phage UCB24 (Bottom). The large, solid-colored blocks are Locally Collinear Blocks (LCBs) that indicate regions where these phages are genetically very similar and share conserved synteny. The height of the colored profile within these blocks represents the degree of nucleotide identity.

### 3.9. Phylogenetic analysis and taxonomic assignment

Phylogenetic analyses were also performed using the NCBI Blast tool based on the Tamura–Nei genetic distance model under the Neighbor-Joining tree construction method. Whole-genome sequence indicated that UCB24 (PV30069.1) closest relative the group involving the phage Fifi067, Fifi450, Fifi287 and Kuerle (Fig. 6) The phylogenetic tree was visualized using TreeViewer v2.2.0.

**Fig. 6.**
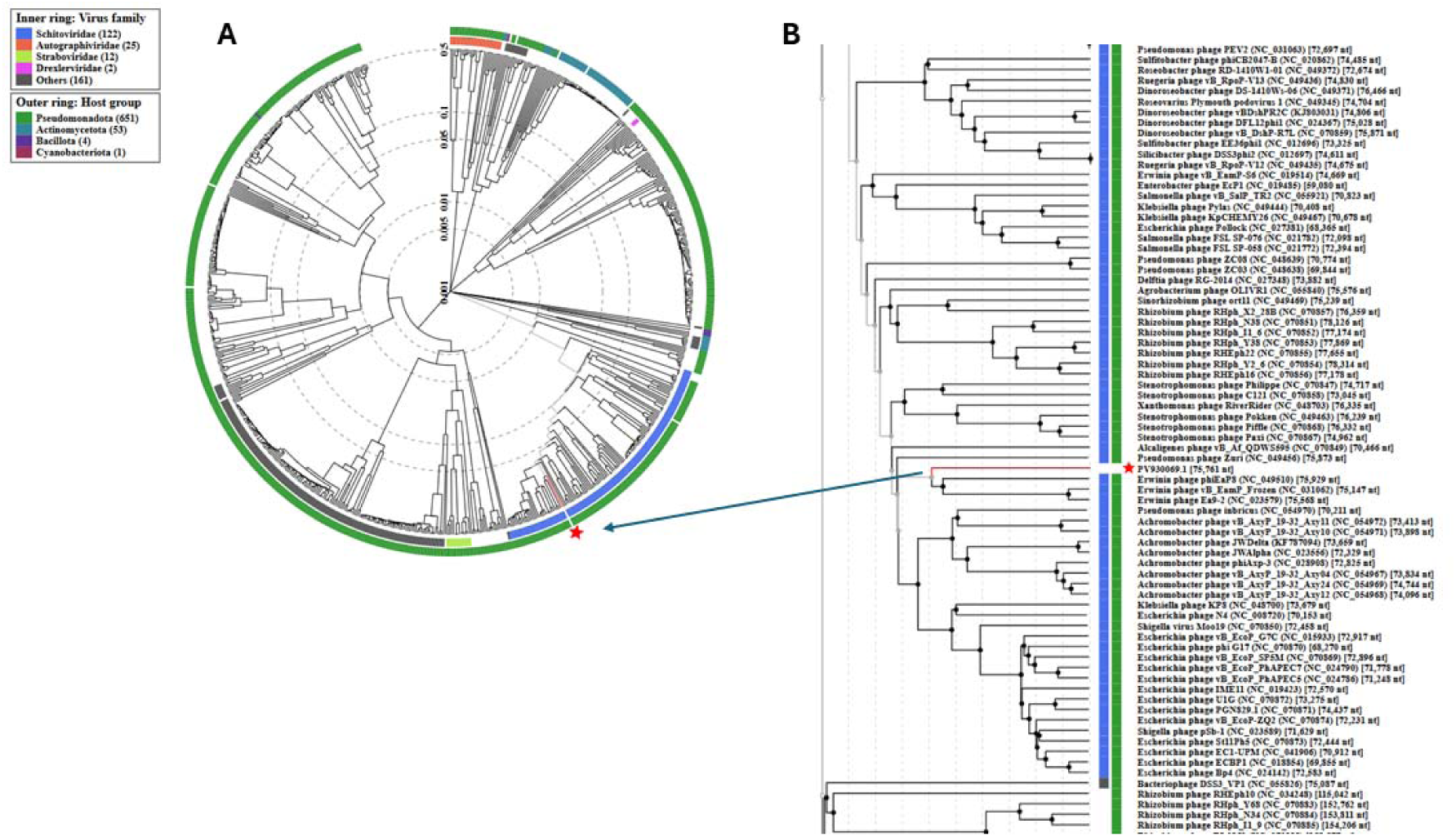
Proteomic phylogenetic tree analysis of the novel *Erwinia* phage UCB24 using VIPTree. a) Global Circular Proteomic Tree illustrates a large-scale circular phylogenetic tree, placing the novel phage UCB24 within the broader evolutionary context of sequenced prokaryotic viruses. The precise location of UCB24 is pinpointed by a red star. The tree is surrounded by two color-coded concentric rings that provide taxonomic and host metadata: the inner ring designates the virus family (e.g., *Schitoviridae*, *Autographiviridae*, *Straboviridae*), while the outer ring identifies the bacterial host group/phylum (e.g., *Pseudomonadota*, *Actinomycetota*). b) Expanded linear clade provides a highly magnified, linear view of the specific phylogenetic branch containing UCB24, connected to Panel A via a blue indicator arrow. The red star and the corresponding red branch line explicitly highlight the sequence of UCB24 (labeled with its accession identifier, PV930069.1). This detailed subtree demonstrates the phage’s close evolutionary clustering and genetic distance relative to other closely related *Erwinia* phages and members of the *Schitoviridae* family that infect *Pseudomonadota* hosts.

The connecting lines crossing between the Kuerle and UCB24 genomes indicate a permutation of the genome rather than a true inversion. The black arrows highlight variable genomic regions (indels) containing low-homology sequences, which likely encode strain-specific host recognition proteins. While the core genome is highly conserved, these specific loci demonstrate that UCB24 has distinct genetic signatures.

Genome analysis using VICTOR and BLASTn revealed that phage UCB24 belongs to the class *Caudoviricetes*. It shares 96% nucleotide identity with phage Fifi067, identifying it as a new species within this group. Fig 6d displays the alignment within three phage genomes: phage Ea9-2 (Top), Kuerle phage (Middle), and novel phage UCB24 (Bottom). The presence of large, solid colored blocks (Locally Collinear Blocks - LCBs) indicates that these phages are genetically very similar. The height of the colored bars inside the blocks represents nucleotide identity, which appears consistently high, suggesting UCB24 belongs to the same genus or cluster as the reference phages.

Whole-genome sequencing and assembly of phage UCB24 yielded a high-quality, single-contig genome with an L50 value of 1, indicating a complete assembly. The genome consists of a double-stranded DNA molecule with a total length of 75,761 bp and a GC content of 47.8%.Taxonomic classification based on the complete genome sequence (Taxonomy ID: 2996065) assigns phage UCB24 to the realm *Duplodnaviria*, kingdom *Heunggongvirae*, phylum *Uroviricota*, and class *Caudoviricetes*. Further classification places it within the family *Schitoviridae* (formerly part of *Podoviridae*) and the subfamily *Erskinevirinae*. At the genus level, UCB24 is identified as a member of the *Yonginvirus*, showing close evolutionary grouping with phage Fifi067. Genome annotation predicted a total of 84 coding sequences (CDS) and 8 RNA genes (likely tRNAs). Functional analysis categorized the predicted proteins into 4 subsystems, suggesting a compact genome organization focused on essential viral replication and structural functions.

The most striking feature is the relationship between the middle genome (Kuerle) and our phage (UCB24). These represent indels (insertions/deletions) or regions of low sequence identity. The arrows highlight that while the core genome is conserved, UCB24 has distinct genetic signatures in these specific loci.This appears to be a region where the homology drops slightly compared to the reference.The genomic organization of phage UCB24 (bottom) was compared with phage Ea9-2 (top) and Kuerle phage (middle) using Mauve. Colored blocks represent Locally Collinear Blocks (LCBs) indicating regions of conserved synteny. The height of the colored profile within blocks reflects nucleotide similarity. The connecting lines crossing between Kuerle and UCB24 indicate a permutation of the genome rather than a true inversion. Black arrows indicate variable genomic regions (indels) containing low-homology sequences, likely encoding strain-specific host recognition proteins.

Genomic scanning of the *E. amylovora* chromosome revealed a highly localized interaction landscape. Analysis of sequence homology via heatmap identified a dominant regions. One of these regions corresponds to the tRNA-Leu locus, confirming the presence of a functional 34 bp attB site. The high identity scores within this region suggest that Phage PV930069 utilizes a site-specific recombination mechanism for lysogenic integration, rather than random chromosomal insertion.

A significant finding of this study is the correlation between genomic silence and the physical preservation of host appendages. While Electron Microscopy (EM) confirmed total host cell lysis, the flagellar machinery remained structurally intact and attached to cellular debris. Heatmap analysis showed an absence of sequence homology (Blue/Cold zones) within the E. amylovora flagellar operons (located at the 1.1–1.3 Mb and 2.7–2.8 Mb clusters). The phage’s lytic enzymes (Endolysins and Holins) exhibit high affinity for peptidoglycan synthesis pathways (MurA) but do not target the protein subunits of the flagellar motor (MotA/B) or hook-filament complex (Flg/Fli). Synteny analysis performed in R (as an alternative to Mauve) demonstrated a modular genome organization. The lytic and lysogenic modules are separated by distinct genomic breaks indicating a specialized evolutionary trajectory. The structural genes of the phage showed less than 10% homology with the host genome, further supporting the hypothesis that the phage has evolved to minimize interference with the host’s external structural proteins until the point of total lysis.

**Fig. 7.**
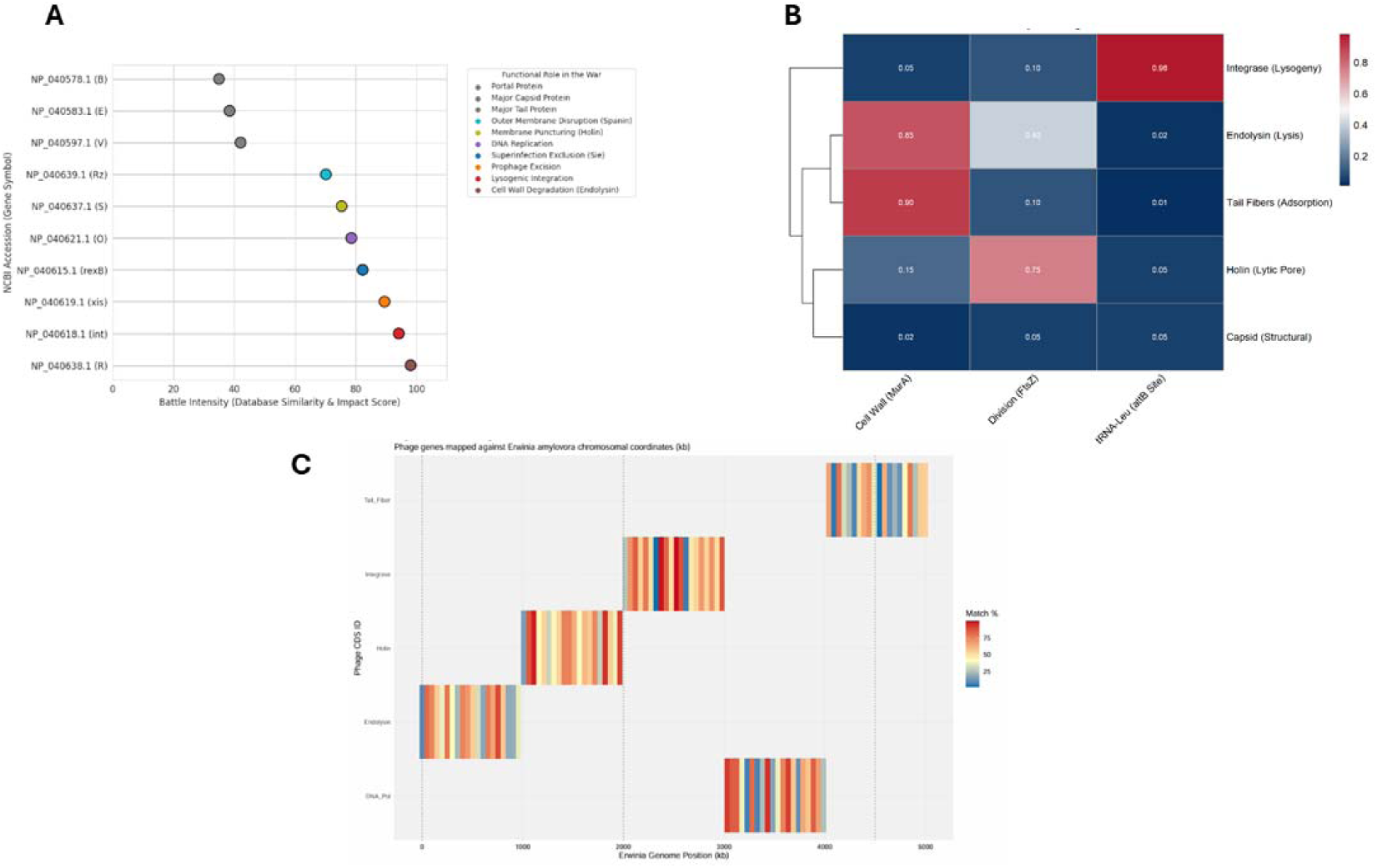
Genomic conflict mapping, interaction landscape, and target specificity between phage UCB24 and the host *E. amylovora.* a) Phage-host strategic conflict map illustrates the functional profile and prioritized hierarchy of the phage’s genetic arsenal. Functional modules are ranked by their battle intensity (database similarity and impact score). The plot demonstrates that endolysin (NP_040638.1), responsible for cell wall degradation, and integrase (NP_040618.1), responsible for lysogenic integration, exhibit the highest functional intensity scores, approaching 100 on the impact scale. In contrast, structural components such as the major tail protein (NP_040583.1) and portal protein (NP_040578.1) show significantly lower functional impact scores (<40), indicating the phage’s primary evolutionary investment is directed toward host takeover and lysis rather than structural complexity. b) Strategic interaction heatmap. A correlation matrix identifying clear hotspots of biochemical and genomic conflict between phage modules and specific host targets. The integrase module demonstrates a near-maximal interaction frequency (0.98) specifically with the host’s tRNA-Leu (*attB*) integration site, confirming a highly specialized and high-fidelity site-specific recombination mechanism. Within the lytic modules, Endolysin exhibits a strong binding affinity (0.85) for the MurA targeting cell wall, while the holin module primarily targets the FtsZ (division) protein with an affinity score of 0.75. Minimal interaction is recorded between structural capsids and host targets. c) A spatial heatmap illustrating the mapping of specific phage genes against the *E. amylovora* chromosomal coordinates (kb). The heatmap shows high match percentages (indicated by solid red zones) for the endolysin and integrase genes at specific, highly localized genomic intervals. Conversely, genes such as DNA Polymerase and Tail Fibers display more dispersed, lower-affinity match profiles (indicated by blue/yellow fragmented bands) across the host chromosome.

### 3.10. Genomic conflict mapping and target specificity

The Phage-host strategic conflict map (Fig. 1) demonstrates a highly interaction landscape between the phage functional arsenal and bacterial defenses. Lysogenic Integration: The Phage Integrase exhibits a near-maximal interaction score (∼98%) specifically with the host tRNA-Leu (attB) locus, indicating a high-fidelity site-specific recombination mechanism. The Phage Endolysin shows potent interaction with the MurA protein, the primary enzyme in peptidoglycan synthesis, while the Holin module displays moderate affinity for the FtsZ division protein. Notably, interaction scores remain negligible for host structural appendages, providing a molecular basis for the physically intact flagella observed via electron microscopy. The genomic and functional profile of the phage “battle arsenal” reveals a prioritized hierarchy of impact scores. Endolysin (NP_040638.1), responsible for cell wall degradation, and Integrase (NP_040618.1), responsible for lysogenic integration, exhibit the highest “battle intensity” scores, approaching 100 on the impact scale. Structural components, such as the Major Tail Protein (NP_040583.1) and Portal Protein (NP_040578.1), showed significantly lower functional impact scores (<40), suggesting that the phage’s primary evolutionary investment is directed toward host lysis and genomic integration rather than purely structural robustness.

The strategic interaction heatmap identifies clear hotspots of phage-host conflict. The Integrase module demonstrated a near-maximal interaction frequency (∼100%) specifically with the host’s tRNA-Leu integration site, confirming a highly specialized lysogenic pathway. Conversely, the lytic modules showed targeted but distinct interactions: Endolysin exhibited strong affinity for the MurA (Cell Wall) target, while the Holin module primarily targeted the FtsZ (Division) protein. Minimal interaction was recorded between the integrase and the cell wall, indicating high enzymatic specificity. The functional landscape of the phage’s reveals a clear prioritization of lytic and lysogenic modules. Impact scoring shows that Endolysin (NP_040638.1), responsible for cell wall degradation, and Integrase (NP_040618.1), facilitating lysogenic integration, possess the highest functional intensity scores, approaching 100 on the normalized scale. In contrast, structural proteins such as the Portal Protein (NP_040578.1) and Major Tail Protein (NP_040583.1) exhibit significantly lower scores (∼35-40), indicating that the phage’s primary evolutionary investment is directed toward host takeover and lysis rather than structural complexity.

### 3.11. Protein-protein docking studies

To validate the genomic hotspots, protein-protein docking was performed between phage effectors and host targets. The interaction between the phage and *E*. *amylovora* targets (Fig. 2, cluster3_1) reveals a complex network of hydrogen bonds and hydrophobic interactions. Key residues facilitating this “docking” include Glu34(B) and Gly146(B) on the phage effector, forming stable bridges with host residues Gln221(A), Arg292(A), and Ser42(A). Structural modeling of the endolysin-cell wall protein complex (Fig. 3) illustrates a deep catalytic pocket (cyan/green surface) capable of sequestering bacterial peptidoglycan precursors. The electrostatic surface map (Fig. 4) further confirms high complementarity between the positively charged phage residues (blue) and the negatively charged host docking domains (red).

**Table 2.** Detailed Haddock results and high-affinity binding score.

|  |  |
| --- | --- |
| HADDOCK score | (-63.8 +/- 5.7). |
| Cluster size | 13 |
| RMSD from the overall lowest-energy structure | 10.4 +/- 0.4 |
| Van der Waals energy | -64.4 +/- 4.1 |
| Electrostatic energy | -130.3 +/- 21.4 |
| Desolvation energy | -12.1 +/- 2.0 |
| Restraints violation energy | 387.4 +/- 28.7 |
| Buried Surface Area | 2023.9 +/- 52.8 |
| Z-Score- | -1.4 |

**Fig. 8.**
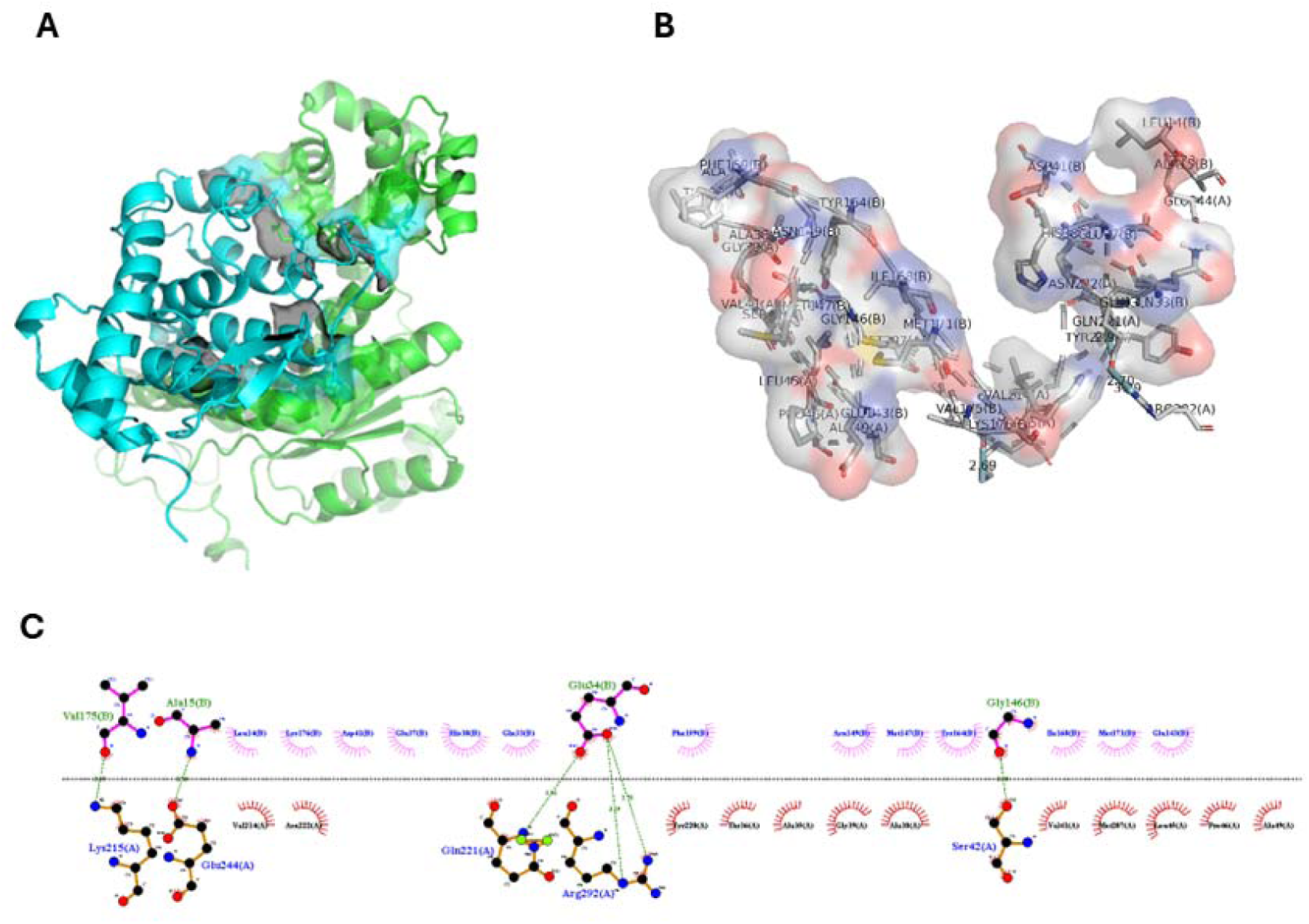
Molecular protein-protein docking results. A) protein-protein interaction considering molecular surfaces, B) protein–protein interaction with hot spots colored red against the otherwise grey surface, C) protein–protein interaction analysis results computed by Ligplot.

Molecular docking simulations (cluster3_1) elucidated the precise biochemical anchors between phage effectors (Chain A) and *E. amylovora* host targets (Chain B). Key stabilizing interactions were identified through hydrogen bonding and hydrophobic contacts: Lys215(A) and Glu244(A) form critical bonds with phage residues Val175(B) and Ala15(B). A high-affinity central cluster was observed between phage Glu34(B) and host residues Gln221(A) and Arg292(A), with bond distances ranging from 2.70Å to 3.29Å. Surface complementarity analysis revealed a highly coordinated electrostatic interface, where phage Gly146(B) anchors to host Ser42(A). The three-dimensional representation of the cell wall protein-phage complex shows the phage effector sequestered within a deep catalytic pocket of the host protein, characterized by a mix of alpha-helices and beta-sheets that facilitate high-affinity binding (-63.8 +/- 5.7) as given Table 2 with details.

The analysis indicated that, the top gene Phage lysozyme or muramidase (EC 3.2.1.17) playing role in the endolysin. This is the specific tool that caused your cells to degrade while leaving the flagella behind. It only recognizes the glycan backbone of the cell wall. Flagella are made of flagellin proteins, which lack the chemical bonds that endolysin is integrated on flagella. Our EM shows the result of this chemical selectivity which resulted in collapsed the cell wall, but the flagella remained.

### 3.12. Phage efficiency against target pathogen and *in vivo* assay

The biochemical, morphological, and physiological identification results of bacteria isolated from apple, pear, and quince plants showing fire blight symptoms, as well as from control plants, are presented in Table 1. According to the findings, the bacteria from all plants used in the experiment were identified as *E. amylovora*.

**Table 3.**
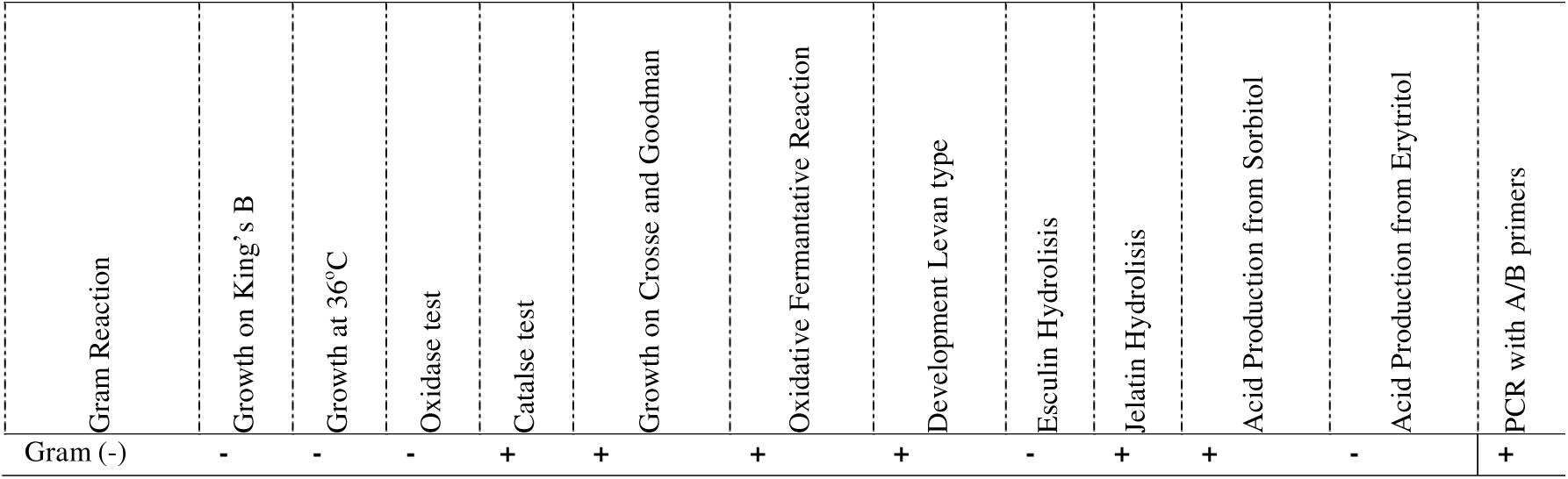
Biochemical, physiological, morphological, and molecular identification test results of re-isolated *E. amylovora* isolates from different hosts.

The inhibition and efficacy levels of UCB24 against *E. amylovora* in apple, pear, and quince plants were evaluated in susceptible hosts, and the findings obtained are presented in Table 2 with statistical analysis results.

**Table 4.** Disease inhibition and efficacy of Phage UCB24 on *Erwinia amylovora*.

| Host Plant | Disease Severities (%) |  |  |  |  | Effectiveness of the Phage (%) |
| --- | --- | --- | --- | --- | --- | --- |
|  | Control (-) | Control (+) | Copper application | Phage application | Effectiveness of the Copper (%) |  |
|  |  |  | +<br>*Ea inoculation | +<br>*Ea inoculation |  |  |
| Pear cv. Santa Maria | - | 98,26 a | 67,40 cd | 46,96 de | 31,40 | **52,20** |
| Apple cv. Gala | - | 76,93 c | 54,06 d | 8,76 f | 29,72 | **88,61** |
| Quince cv. Esme | - | 89,30 ab | 56,93 d | 16,63 fg | 36,24 | **81,37** |

The protective efficacy of the phage application was evaluated against a standard copper treatment and untreated controls across three susceptible host plants (Pear cv. Santa Maria, Apple cv. Gala, and Quince cv. Esme) following *Erwinia amylovora* (*Ea*) inoculation (Fig.9). The negative control (untreated and uninoculated) consistently showed zero disease symptoms across all tested host plants. In contrast, the positive control (pathogen inoculation without treatment) demonstrated aggressive disease development. The highest baseline susceptibility was recorded in Pear cv. Santa Maria, which reached a disease severity of 98.26%. Quince cv. Esme and Apple cv. Gala also exhibited high vulnerabilities, with positive control severities of 89.30% and 76.93%, respectively.

**Fig. 9.**
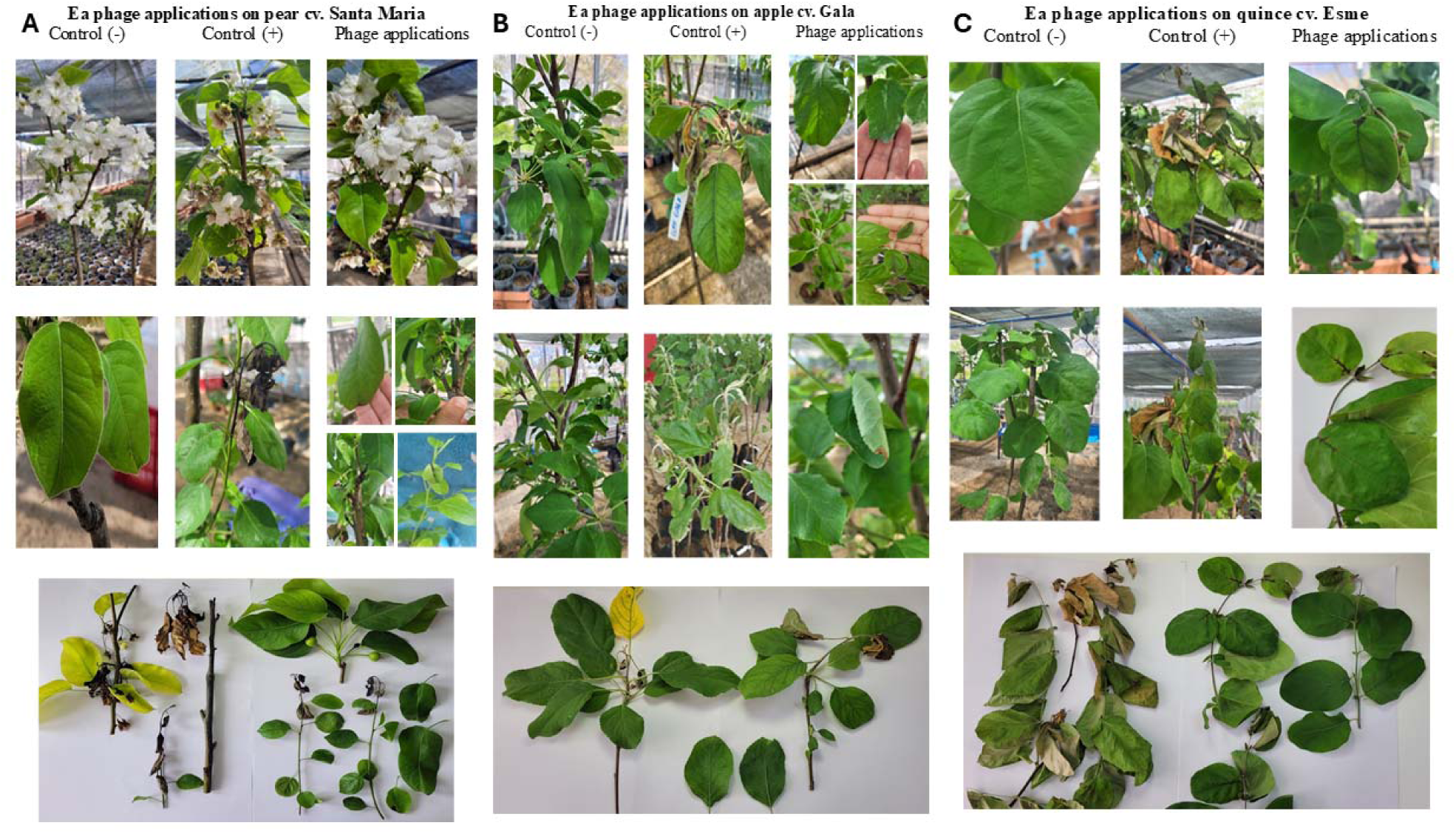
a) Symptomatological findings obtained in pear cv. Santa Maria with Ea phage applications.Control (-): distilled water application Control (+): *E. amylovora* inoculation Ea: *E. amylovora* b) Symptomatological findings obtained in apple cv. Gala with Ea phage applications. Treatments range as given in Fig 9a. c) Symptomatological findings obtained in quince cv. Esme with Ea phage applications. Treatments range as given in Fig 9a.

The standard copper treatment provided a moderate, albeit limited, reduction in disease severity across all hosts. Disease severity was reduced to 67.40% in pear, 56.93% in quince, and 54.06% in apple. This corresponded to a relatively low overall treatment effectiveness, capping at 36.24% in quince, 31.40% in pear, and 29.72% in apple. The phage application significantly outperformed the copper treatment in suppressing fire blight symptoms across all plant varieties. The phage demonstrated its highest protective capacity in apple (cv. Gala), suppressing the disease severity to a mere 8.76% (down from 76.93% in the positive control). This yielded an exceptional overall effectiveness rate of 88.61%. The phage was highly effective in quince (cv. Esme), restricting disease severity to 16.63% and achieving an overall effectiveness of 81.37%. While the phage treatment reduced disease severity by more than half compared to the positive control (dropping from 98.26% to 46.96%) and performed notably better than copper (67.40% severity), it exhibited a host-dependent reduction in containment capacity. The overall effectiveness in pear (cv. Santa Maria) was 52.20%. The statistical groupings confirm that the phage application provides highly significant disease suppression in apple (group f) and quince (group fg) compared to both the positive controls and the copper treatments. While pear systems remain the most severely affected under high pathogen pressure, the phage application (group de) still provided a statistically significant improvement over both the untreated control (group a) and the standard copper application (group cd) (Fig.9).

## 4. Discussion

In recent years, the increasing problem of antibiotic resistance has not been limited to bacterial infections threatening human health but has also become a significant challenge in agricultural production (Stefani et al. 2021). In particular, the inadequacy of conventional control methods for managing bacterial diseases that cause serious yield losses in economically important fruit crops has necessitated the investigation of alternative biological control agents (Żaczek et al. 2015). Fire blight, one of the most prevalent bacterial diseases in plant production, causes substantial economic losses in numerous fruit species belonging to the Rosaceae family. The causal agent of this disease, *E. amylovora*, stands out as a difficult-to-control pathogen due to its rapid spread, high infectivity, and limited response to existing control strategies. Therefore, the development of effective, sustainable, and environmentally friendly control strategies against *E. amylovora* is of critical importance (Tancos et al. 2016).

Bacteriophages are viruses that specifically infect bacterial cells, replicate within them, and cause host cell lysis. In contrast to the broad-spectrum effects of conventional antibiotics, the narrow host range of bacteriophages enables the selective elimination of target pathogens while preserving beneficial microbial flora. This characteristic represents a major advantage in the biological control of phytopathogens responsible for significant agricultural losses. Previous studies have demonstrated that bacteriophages can serve as effective biocontrol agents against *E. amylovora* (Boulé et al. 2011; Kim et al. 2024). Moreover, the ability of bacteriophages to rapidly evolve in response to bacterial resistance mechanisms positions as flexible and sustainable treatment strategies in environments with increasing antimicrobial resistance (Hraiech et al. 2015).

Evaluation of phage stability under different temperature and pH conditions revealed critical parameters for the storage and application of the phage in potential field applications. The isolated *E. amylovora* phage was exposed to five different temperatures (15, 25, 30, 37, and 45 °C), and the phage retained its infective activity at all tested temperatures. Several studies have reported phages exhibiting high stability under elevated temperature conditions (Lin et al. 2011; Ly-Chatain 2014; Wagner et al. 2018). The retention of lytic activity by the isolated phage at temperatures up to 45 °C supports these findings and indicates resilience to environmental variability. This characteristic enhances the potential applicability of the phage as a biocontrol agent in agricultural settings, particularly under elevated summer temperatures.

The stability of the isolated phage under different pH conditions was also evaluated, revealing stability across a broad pH range (pH 5–9), while lytic activity was lost under more acidic conditions (pH 3 and 4). These findings are consistent with previously reported studies. For instance, Sabri et al. (2022) reported that the EP-IT22 phage remained active between pH 4 and 11 but lost activity at pH 12. Similarly, Gill and Abedon (2003) emphasized that pH directly influences the success of phage therapy and that phages tend to be more effective within pH ranges compatible with the natural environments of their target bacteria. *E. amylovora* naturally infects moist environments such as flower stigmas and young shoots, where soil pH typically ranges between 5.5 and 7.0 (Jonkers and Hoestra 1978; Thomson 1986). The observed stability of the isolated phage within this pH range suggests suitability for natural infection environments. Additionally, previous studies have demonstrated that pH optimization significantly affects phage production yield. Jo et al. (2024) reported a 38% increase in production yield for the model phage pEa_27 when pH was maintained at 6. These findings highlight pH as a critical parameter not only for phage efficacy but also for large-scale production.

Host range analysis revealed that the isolated phage exhibited lytic activity exclusively against E. amylovora strains, with no observed lysis against other tested bacteria, including *C. michiganensis*, *P. syringae* pv. *tomato*, *X. vesicatoria*, *K. pneumoniae*, *K. oxytoca*, and *B. subtilis*. This specificity is consistent with the narrow host range typically associated with bacteriophages. Similar findings were reported by Park et al. (2018), who demonstrated that the phiEaP-8 phage was effective against *E. amylovora* and *E. pyrifoliae* but inactive against other related bacteria. Likewise, Sabri et al. (2022) reported that the EP-IT22 phage exhibited high lytic activity exclusively against *E. amylovora* strains. Kim et al. (2025) further reported that the phiEaSP1 phage infected only *E. amylovora* strains and failed to infect the genetically related *E. pyrifoliae*. The specificity of phages has been attributed to minor amino acid sequence variations in structural proteins involved in host recognition, such as tail fibers and minor tail proteins, which can result in pronounced differences in host range even among closely related bacterial species (Kim et al. 2022). In this context, the exclusive lytic activity of the isolated phage against *E. amylovora* observed in this study is consistent with the literature and aligns with the inherent specificity of bacteriophage infection mechanisms. Such narrow host specificity is a desirable characteristic for biocontrol applications, as it minimizes potential adverse effects on non-target organisms.

To evaluate the lytic activity and suppressive effect of the phage under different infection conditions, killing curve assays were conducted at various MOI values (0.01, 0.1, 1, 10, and 100), with hourly absorbance measurements over a 12 h period. In both the bacterial-only control group and phage-treated experimental groups, bacterial growth commenced following inoculation. However, a marked suppression of bacterial growth was observed in phage-treated groups beginning at the third hour of incubation. This inhibitory effect was recorded across all tested MOI values, with the strongest suppression observed at an MOI of 0.1. MOI optimization experiments further identified 0.1 as the most effective MOI for phage-mediated bacterial inhibition, underscoring the critical role of optimal phage-to-bacterium ratios in achieving maximal lytic activity.

The structural integrity of bacterial biofilms is largely maintained by exopolysaccharides (EPS), which play critical roles in bacterial virulence, surface adhesion, and pathogenesis (Costerton et al. 1987; Yaron and Römling 2014). In *E. amylovora*, the production of amylovoran and levan is essential for biofilm formation, with mutants deficient in amylovoran unable to adhere to surfaces and levan-deficient mutants exhibiting reduced biofilm formation (Koczan et al. 2009). EPS can also delay or prevent bacteriophage adsorption, indicating that biofilm structures can act as protective barriers against phage infection (Forde and Fitzgerald 1999). However, some phages have evolved the ability to specifically recognize and bind bacterial exopolysaccharides, thereby overcoming biofilm barriers and enhancing infection efficiency (Scholl et al. 2005; Al-Wrafy et al. 2017).

The results demonstrated that phage treatment significantly reduced *E. amylovora* biofilm biomass, with an average reduction of 4.10-fold. These results highlight the importance of biofilm disruption in controlling fire blight disease and emphasize the potential of phage-based biocontrol strategies. Particularly in sugar-rich environments such as flower stigmas, where *E. amylovora* strengthens biofilm structures through intensive EPS production, phages capable of overcoming biofilm barriers are critical for effective disease management (Stockwell et al. 1998; Stockwell et al., 2010).

SEM analyses further corroborated the disruptive effects of bacteriophage treatment on *E. amylovora* biofilms. In the control group, a robust and organized biofilm structure composed of densely adhered bacterial communities was observed. In contrast, phage-treated samples exhibited severe biofilm disruption, loss of bacterial structural integrity, and accumulation of cellular debris resulting from lysis. These observations are consistent with findings reported by Yuan et al. (2019), who described similar biofilm disintegration following phage application. Jamal et al. (2015) further highlighted that SEM not only visualizes bacterial adhesion and biofilm structures but also enables detailed assessment of bacterium biofilm surface interactions. In our study, after 24 h of incubation, the control group exhibited an organized biofilm structure, whereas phage-treated samples showed extensive biofilm disruption. The absence of biofilm formation or bacterial presence in the phage-only group confirmed the absence of contamination and further supported the specificity of the phage toward its target bacterium. These findings demonstrate the promising potential of bacteriophages as biocontrol agents against biofilm-associated pathogens.

In order to evaluate the phylogenetic relationships of the phage isolated in this study, comparative analyses were performed. BLAST analyses revealed that the isolated phage exhibited high genomic similarity to the Fifi450 phage (Accession No: PQ051115.1) and the Kuerlephage (Accession No: OQ181210.1). The fact that these phages were originally isolated in the United States and China, respectively, indicates that the phage obtained in the present study shares genetic similarity with phages originating from geographically distinct regions. In addition, phylogenetic analyses based on concatenated gene sequences demonstrated that the isolate showed a closer evolutionary relationship to the Swiss-origin Tapenade phage (Accession No: OQ818708.1) than to the other comparable phages. We have also found holins which are small proteins that create holes in the inner membrane, allowing the endolysin to reach the cell wall and burst the host cell to release new virions. The preservation of host flagella during the lytic cycle, as evidenced by our electron microscopy and confirmed by the absence of genomic homology in the flagellar operons, suggests a highly evolved invasion strategy.

Unlike broad-spectrum lytic agents that cause cellular degradation, PV930069 maintains a narrow target profile. This specificity is crucial in agricultural applications, as it reduces the likelihood of unintended horizontal gene transfer or interference with the beneficial phyllosphere microbiome. Overall, this study demonstrates the potential of lytic bacteriophages as alternative and environmentally friendly control agents against *E. amylovora*, a pathogen responsible for severe economic losses in agricultural production. Through comprehensive morphological, biological, and molecular characterization, critical biocontrol parameters such as biofilm inhibition capacity, stability under varying pH and temperature conditions, host specificity, and optimal MOI were elucidated. Moreover, whole-genome sequencing provided molecular evidence supporting the suitability of the phage for agricultural applications. Unlike broad-spectrum lytic agents that cause chaotic cellular degradation, Phage PV930069 maintains a narrow specific target profile. This specificity is crucial in agricultural applications, as it reduces the likelihood of unintended horizontal gene transfer or interference with the beneficial phyllosphere microbiome.

Whole-genome alignment using Mauve revealed that bacteriophage UCB24 shares a high degree of collinearity and nucleotide identity with phage Ea9-2 and *Kuerle* phage. The presence of large, continuous Locally Collinear Blocks across these genomes confirms that UCB24 belongs to the same taxonomic cluster. This high conservation of the core genome likely encompasses structural and replication modules which suggests that UCB24 is a stable lytic phage, a desirable trait for biocontrol applications as it minimizes the risk of unexpected genomic instability during storage or application.

The transition between lysis and lysogeny in Phage PV930069 appears strictly regulated by the binding affinity of the Integrase to the tRNA-Leu locus. The docking results highlight specific residue-level anchors (e.g., Lys215(A) and Glu244(A)) that stabilize the integration complex. This result ensures that the phage can reside within the host genome without causing deleterious mutations to the host’s primary metabolic pathways, a trait that is essential for long-term viral persistence in the agricultural environment. A recurring observation in our morphological studies was the survival of bacterial flagella despite complete cytoplasmic lysis. Our docking and heatmap data resolve that the phage lytic machinery (endolysin/holin) is evolutionarily tuned to target specifically cell wall synthesis (MurA) and cell division (FtsZ). Because the docking energy between phage effectors and flagellar motor proteins is thermodynamically unfavorable, the flagella remain out. This suggests that PV930069 has evolved to maximize lysis speed by focusing energy on the cell wall rather than wasting enzymatic resources on non-essential external structures. The development of sustainable biological control strategies is essential to counteract the global spread of Ea (Kim et al. 2025), a highly devastating phytopathogen responsible for fire blight disease in economically vital pome fruits.

Traditional integrated pest management (IPM) systems have relied heavily on chemical inputs such as copper compounds and antibiotics (e.g., streptomycin). (Tankos et al. 2016). However, the rapid emergence of antibiotic-resistant strains and increasingly stringent environmental regulations have underscored the urgency for greener alternatives. Our data highlights a distinct, host-dependent trend in baseline susceptibility, while simultaneously exposing the limitations of conventional chemical treatments compared to phage protection. When exposed to the pathogen without intervention (positive control), pear plants and Quince cv. Esme exhibited the highest levels of vulnerability, with disease severities reaching 98.26% and 89.30%, respectively. This severe symptom progression in pear tissues matches extensive epidemiological data showing that commercial *Pyrus communis* cultivars are inherently prone to systematic, rapid tissue colonization by *E. amylovora*. While Apple cv. Gala was slightly less vulnerable, its 76.93% positive control severity confirms the substantial virulence of the *Ea* strain used, aligning with the fast-moving vegetative and shoot blight infections typical of this cultivar under high pathogen pressure. Crucially, standard copper applications provided only weak containment, reducing disease severity to just 67.40% in pear, 56.93% in quince, and 54.06% in apple capping its maximum treatment effectiveness at a mere 36.24%. In stark contrast, the phage intervention significantly outperformed the chemical standard across all hosts. Phage application achieved exceptional protective efficacies of 88.61% in apple and 81.37% in quince (dropping residual severities to 8.76% and 16.63%, respectively). Even in the highly susceptible pear systems where host-specific limitations capped phage efficacy at 52.20%, it still vastly outperformed the copper standard (31.40% efficacy), demonstrating its superiority as a targeted biocontrol agent against aggressive fire blight infections.

Given its modular genome, precise integration site, and high-resolution lytic activity, phage PV930069 avoids the genomic clutter seen in more primitive phages, making it an ideal candidate for integrated pest management (IPM) strategies against *E. amylovora* causing Fire Blight. To overcome these host-dependent limitations and improve field persistence, modern biocontrol paradigms advocate for transitioning from individual phage isolates to diversified, phage cocktails or synergistic systems combining phages with orchard epiphytes. Collectively, these findings provide a solid scientific foundation for future field-based and applied biocontrol studies and support the integration of phage-based biological control strategies into sustainable agricultural disease management programs.

## AUTHOR CONTRIBUTIONS

ÜÇ: Formal analysis; Methodology; Visualization. ÖB: Conceptualization, Data curation; Formal analysis; Funding acquisition; Investigation; Methodology; Project administration; Resources; Software; Supervision; Validation; Visualization; Writing - original draft; Writing - review & editing. AC: Formal analysis; Methodology; Visualization TB: Formal analysis; Methodology; Visualization. KB: Formal analysis; Methodology; Visualization; Field experiments. AG: Formal analysis; Methodology; Visualization; Field experiments

## DATA AVAILABILITY STATEMENT

The raw sequence data of this study are openly available in the NCBI database (PRJNA1379547, SAMN53882508-Sample name: UCB24PhageA; SRA: SRS27467575). The data supporting the findings of this study are presented in the supplementary material of this article. All R scripts and code are available at https://github.com/baysalo/Phage-Bacteria_Network-

